# Exploring the ageing methylome in the model insect, *Nasonia vitripennis*

**DOI:** 10.1101/2023.02.14.528436

**Authors:** K. Brink, C.L. Thomas, A. Jones, T.W. Chan, E. B. Mallon

## Abstract

**Background:** The ageing process is a multifaceted phenomenon marked by the gradual deterioration of cellular and organismal functions, accompanied by an elevated susceptibility to diseases. The intricate interplay between genetic and environmental factors complicates research, particularly in complex mammalian models. In this context, simple invertebrate organisms have been pivotal, but the current models lack detectable DNA methylation limiting the exploration of this critical epigenetic ageing mechanism.

This study introduces Nasonia vitripennis, the jewel wasp, as an innovative invertebrate model for investigating the epigenetics of ageing. Leveraging its advantages as a model organism and possessing a functional DNA methylation system, Nasonia emerges as a valuable addition to ageing research.

**Results:** Whole-genome bisulfite sequencing unveiled dynamic alterations in DNA methylation, with differentially methylated CpGs between distinct time points in both male and female wasps. These changes were associated with numerous genes, enriching for functions related to telomere maintenance, histone methylation, and mRNA catabolic processes. Additionally, other CpGs were found to be variably methylated at each timepoint. Sex-specific effects on epigenetic entropy were observed, indicating differential patterns in the loss of epigenetic stability over time. Constructing an epigenetic clock containing 19 CpGs revealed a robust correlation between epigenetic age and chronological age.

**Conclusions:** Nasonia vitripennis emerges as a promising model for investigating the epigenetics of ageing, shedding light on the intricate dynamics of DNA methylation and their implications for age-related processes. This research not only expands the repertoire of ageing models but also opens avenues for deeper exploration of epigenetic mechanisms in the context of ageing.

## Introduction

Ageing is a complex biological process characterized by a progressive decline in cellular and organismal function, accompanied by an increased susceptibility to diseases. Ageing is influenced by many environmental and genetic components. The effects of these components influence each other making them difficult to investigate, especially in complex mammalian models. Therefore, a large body of ageing research is based on simple invertebrate model organisms [1, 2]. Advantages include easy and cheap to keep in a laboratory, short life span, genetic and molecular tools available and a sequenced genome. However, the current invertebrate models of ageing (Drosophila [3] and *C. elegans* [4]) do not possess detectable DNA methylation.

DNA methylation involves the addition of a methyl group to cytosine residues, predominantly occurring at cytosine-phosphate-guanine (CpG) dinucleotides. Studies have consistently demonstrated dynamic changes in DNA methylation patterns during ageing, influencing gene expression, cellular function, and ultimately contributing to the ageing phenotype [5]. Despite not being found in our current two invertebrate models of ageing, this is not true universally in invertebrates and changes in DNA methylation have been associated with ageing in ants [6], bumblebees [7] honeybees [8] and the crustacean *Daphnia magna* [9]. Although social insects present an almost unique opportunity for an invertebrate model of the effects of social environment on ageing [10], their colony structure prevent them from being easily kept general models of ageing. Although *Daphnia magna* is relatively easy to keep, as a crustacean we can not take advantage of the large amount of research on hymenopteran DNA methylation [11].

We propose the jewel wasp, *Nasonia vitripennis* as a general model for epigenetic ageing as it possesses both the advantages of a model systems mentioned above [12] and has a functional methylation system [13, 14]. DNA methylation has been shown to be vital for *Nasonia* development [15]. It has also been shown to be invovled in a number of *Nasonia* phenotypes including diapause [16] and sex allocation [17]. Ageing in *Nasonia* has been studied with reference to lipogenesis [18], diet [19], ploidy [20], diapause [21] and host species [22]. To our knowledge, the changes in DNA methylation as *Nasonia* ages have never been measured. Here, we establish *Nasonia* as a model to investigate the epigenetics of ageing by first measuring its methylome as it ages.

The omics era has seen a rapid growth in studies measuring these dynamic changes in DNA methylation during ageing. They are usually based on whole genome bisulphite sequencing (WGBS) [23]. Perhaps the most obvious way to search for patterns in this WGBS data are differentially methylated positions (DMPs). These are characterised by changes in average DNA methylation levels across individuals or tissues at single base positions (CpGs). These CpGS either become more methylated as an organism ages (hypermethylation) or less methylated (hypomethylation). These DMPs represent single base pair changes in DNA methylation as the organism ages.

Epigenetic drift is the increased variability in the epigenome found through the course of an individual’s life caused by the accumulation of mistakes in preserving epigenetic patterns. Epigenetic drift leads to a decrease in the body’s ability to maintain homeostasis [24]. This can be measured in at least two ways. Variably methylated positions (VMPs) rather than comparing the average methylation at a CpG over time, measures how variable the methylation at a site is as organisms age. An alternative measure of epigenetic drift is Shannon’s entropy, i.e. the loss of information in the whole epigenome over time [25]. An increase in entropy means the epigenome is becoming less predictable, that is more variable over time.

An epigenetic clock is an emergent property of the DNA methylation status of a large number of genes, calculated using supervised machine learning methods [25, 26, 27]. There is evidence epigenetic age mirrors true biological age and its associated morbidity and mortality better than chronological age [5]. However, their utility as measures of changes in biological age for clinical interventions is limited as their mechanistic basis is not understood [28]. In this paper, we measure chronological ageing and changes in the methylome using whole genome bisulfite sequencing (WGBS) (Figure 1) in order to discover if *Nasonia vitripennis*, unlike the other two invertebrate models of ageing possesses an ageing methylome.

**Figure 1:**
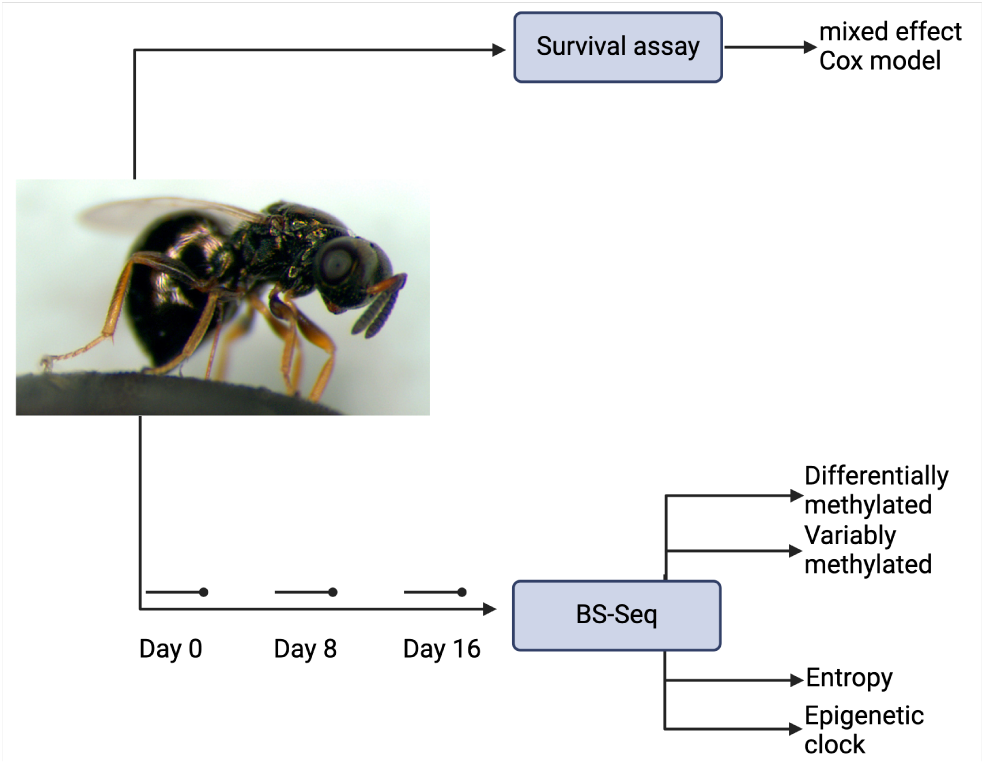
Schematic overview of experimental design with study organism, *Nasonia vitripennis*. Image credit: M.E. Clark public domain.

## Methods

### Life span

*Nasonia* were of the *Nasonia vitripennis* species from the Leicester strain which has been kept for over eight years and originated from the AsymC strain. Wild-type wasps were maintained at 25°C, 40% humidity in a 12-h dark/light cycle. Adults, within 24 hours of eclosion, were placed in tubes of ten single-sex individuals. They were fed 20% sucrose *ad libitum*, refreshed daily. These were checked every day for survival. 70 females and 67 males were used. A mixed effect Cox model treating tube as a random effect was implemented using the survival package (v.3.4) [29] and coxme package (v.2.2) [30] in R 4.2.2 [31].

### DNA extraction

Wasps were collected within twenty four hours of eclosion. Some females may have been mated as they were allowed to mix with males for upto the first fifteen hours after eclosion. Wasps were then collected under light CO_2_ anesthesia and placed into single-sex vials containing ten individuals. They were then provided with filter paper soaked in 20% sucrose which was changed daily. Day zero wasps were collected at the end of the first twenty four hours after eclosion then samples on the eighth and sixteenth day. Thirty wasps of each sex were used for each time-point. Ten wasps from the same sex were pooled for each replicate, creating three replicates for each sex at each time point. Therefore we had three replicates for males and three replicates for females at each of day zero, eight and sixteen after eclosion. Wasps were immediately frozen in liquid nitrogen and stored at minus 80°C freezer for sequencing. DNA was extracted using Qiagen’s DNAeasy Blood and Tissue kit. DNA quality was assessed by NanoDrop 2000 spectrophotometer (Thermo Scientific), 1% agarose gel and Qubit (dsDNA BR Assay, ThermoFisher).

### Whole genome bisulfite sequencing

WGBS sequencing was carried out by BGI Tech Solution Co., Ltd.(Hong Kong). A 1% unmethylated lambda spike was included in each sample in order to assess bisulfite conversion rates. For WGBS samples, library quality was checked with FastQC (v.0.11.5; 32). One day eight male library appeared to be mislabelled by the sequencing company. This was not included in further analysis to prevent any uncertainty. Therefore for the BS-seq analysis we had three replicates for males and three replicates for females at each of day zero and sixteen after eclosion but three replicates for males and two replicates for females at day eight. Paired-end reads were aligned to the *Nasonia vitripennis* reference genome (Nvit PSR1.1, Refseq accession no. GCA 009193385.1, 33) using the Bowtie 2 aligner (v.2.2.9; 34) within the Bismark software (v.0.18.1; 35) under standard parameters. Samples sequenced across multiple files were merged using samtools (v.1.9; 36). Files were deduplicated using Bismark, and methylation counts were extracted in different contexts using the bismark methylation extractor command (v.0.18.1; 35). Destranding was carried out using the coverage2cytosine script from Bismark using the merge CpG command to increase coverage by pooling the top and bottom strand into a single CpG [35]. Reads were also aligned to the unmethylated lambda reference genome to calculate the error rate of the C–T conversion (Refseq accession no. GCF 000840245.1).

### Differential methylation analysis

Output from the coverage2cytosine script was then inputted into the R package methylKit (v.3.14; 37) where files were filtered and normalised based on coverage, removing sites with abnormally high coverage (greater than 99% percentile) or with a coverage less than ten in each sample.

A binomial test was then applied to the filtered CpG sites where the lambda conversion rate was used as the probability of successes and a false discovery rate (FDR) of p < 0.05 [38]. As the majority of sites in the *Nasonia* genome show zero methylation, only CpGs which were methylated in at least one sample were retained. On these methylated CpGs, differential methylation analysis was performed using the calculateDiffMeth command in methylKit, which implements a logistic regression model. Differentially methylated CpG sites were classed as having a minimum difference of > 15% methylation and a q-value < 0.05. Differential methylation analyses were performed across age in each sex.

Genes were classed as differentially methylated if they contained at least two differentially methylated CpG and a minimum weighted methylation difference of 15% across the entire feature [39]. Weighted methylation level is classed as the total number of methylated cytosines (C) within a region (i), divided by the total coverage of that region [39].

### Variable methylation analysis

Variable methylation analysis was carried out on the methylated cytosines. A beta regression model was applied, with methylation proportion as the response variable, and chronological age and sex as predictor variables for each cytosine. The beta regression was implemented using the betareg function from the R package betareg (version 3.1.4) [40].

To identify Variable Methylated Positions (VMPs), the Breusch-Pagan test for Heteroscedasticity was performed using the bptest function in the R package lmtest (version 0.9.40) [41]. Multiple testing corrections were applied using the Holm-Bonferroni method. CpG sites demonstrating significant heteroscedasticity were classified as age-related VMPs.

### Epigenetic drift

Entropy is calculated as;

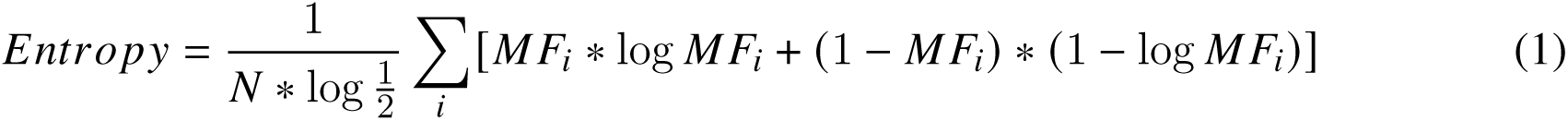

with *MF_i_* the fraction of methylation on a given CpG and N the total number of CpGs measured (5290 significantly differentially methylated CpGs).

The effects of chronological age and sex on entropy were analysed using a beta regression using the betareg package in R [42]. Post-hoc tests were carried out using the emmeans package in R [43].

### Elastic net regression

Chronological age was regressed against the 5290 age significant CpGs’ beta values using an elastic net regression with a 3-fold cross validation implemented in the glmnet R package [44]. The reported accuracy of the epigenetic clock is expected to be overly optimistic since the regression model used cytosines that relate to age in the entire data set. The elastic net regression identified 19 CpGs that predict age. The epigenetic age of each replicate is predicted based on these CpGs methylation state. This epigenetic age was correlated with chronological age using a spearman’s rank correlation. Age acceleration was calculated as the residual from regressing chronological age against epigenetic age [27].

### Gene ontology enrichment

Gene ontology (GO) terms for *N. vitripennis* were taken from the ensembl metazoa database [45]. GO enrichment analysis was carried out using the hypergeometric test with Benjamini-Hochberg [46] multiple-testing correction, q <0.05. GO terms from variably methylated genes and epigenetic clock genes were tested against a GO term database made from the GO terms associated with all methylated genes. Genes were determined as methylated if they had a mean weighted methylation level taken across all replicates greater than the lambda spike weighted methylation level of 0.05 in any of the samples within each comparison. Genes which were consistently hyper or hypo methylated in males or females were tested against all differentially methylated genes in any comparison. REVIGO [47] was used to generate treemaps.

## Results

### Survivourship analysis

Males and females showed different patterns of life expectancy (Cox mixed-effects model: Hazard ratio for females = 2.48 (standard error of coefficient = 0.233), z = 3.89, p = 9.8 × 10^−5^), with females’ mean life expectancy being 29 days and males’ being 17 days, see Figure 2a. This was greater than the 16.6 days for females and 10.7 days for males previously found for sucrose-fed individuals [48].

**Figure 2:**
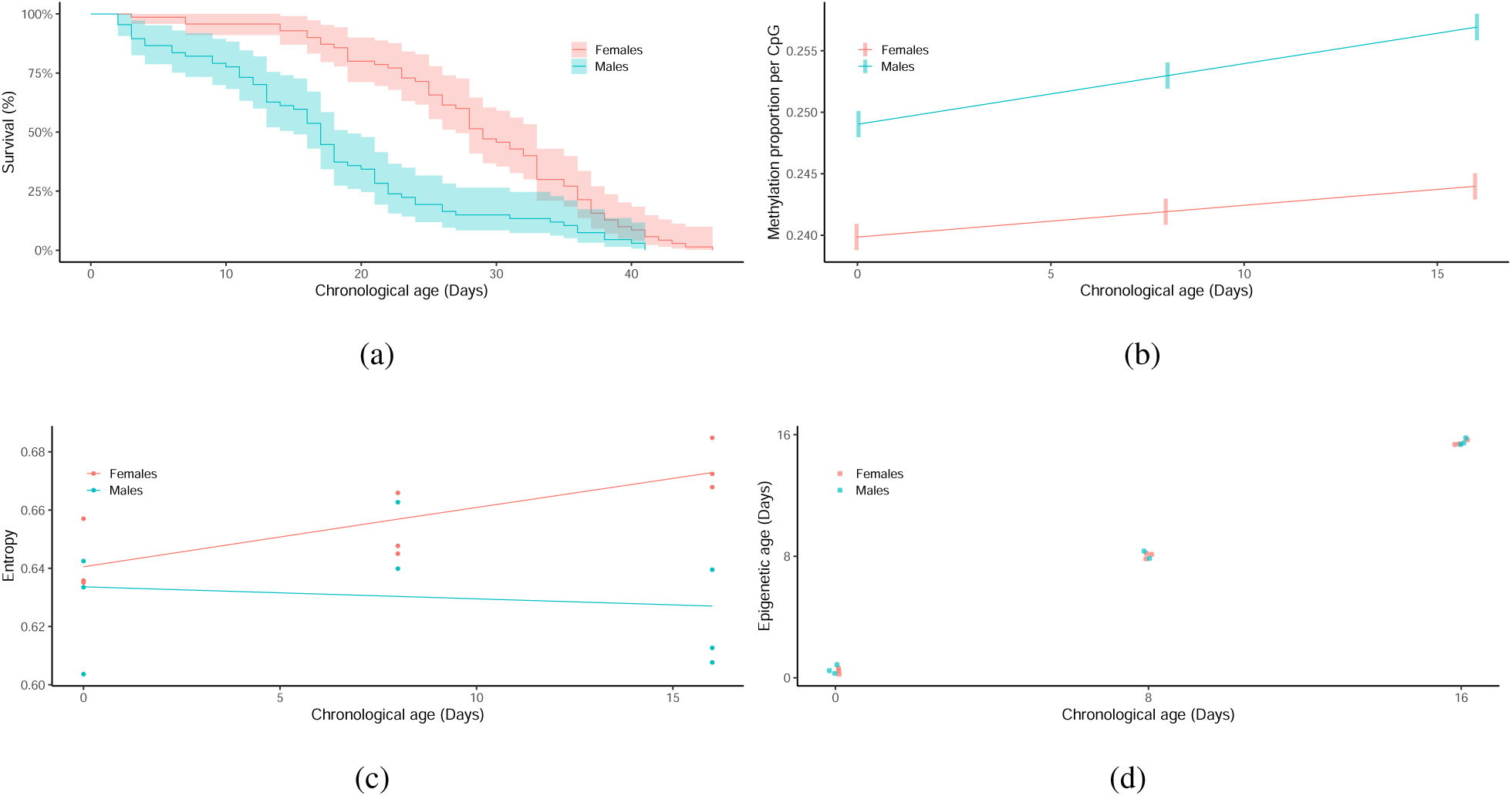
(a) Kaplan-Meier survival curves for female (n = 70) and male (n = 67) *Nasonia vitripennis* adults. Shaded areas represent the 95% confidence intervals. (b) Estimated marginal means of methylation proportion per methylated CpG against chronological age. Shaded bars are 95% confidence intervals (c) Scatterplots of epigenetic entropy based on the 5290 age-related differentially methylated CpGs over time for male and female *Nasonia*. Each point is a single WGBS library made up of ten individuals. Lines represent the beta regression predictions (d) Scatterplot of epigenetic age versus chronological age of male (n=8) and female (n = 9) *Nasonia* samples. Each sample is made up of the whole bodies of ten individuals.

### Differential methylation analysis

5290 CpGs were found to be significantly differentially methylated between at least two time points in males or females. Of these 48% were hypomethylated and 52% hypermethylated. These CpGs were found in 2295 annotated genes (Supplemental Table S1). As males aged, 280 CpGs were hypermethylated consistently, corresponding to 114 annotated genes (Supplemental Table S2). 218 CpGs were consistently hypomethylated as males age (83 genes - Supplemental Table S3). For females, 58 CpGs (26 genes - Supplemental Table S4) were consistently hypermethylated and 34 CpGs (14 genes - Supplemental table S5) hypomethylated. There was no overlap between these groups of CpGs.

In order to explore the function of these genes, we carried out a GO enrichment analysis of the four groups of consistently methylated genes compared to all significantly differentially methylated genes. Genes which are consistently hypermethylated as males age are enriched for telomere function including “*regulation of telomere maintenance*” (GO:0032204), “*telomere organization*” (GO:0032200) and “*meiotic attachment of telomere to nuclear envelope*” (GO:0070197) (Supplemental Figures S1). Genes which are consistently hypomethylated as males age are enriched for GO terms associated with histone methylation including “*regulation of histone methylation*” (GO:0031060) and “*histone H3-K4 trimethylation* (GO:0080182). “*response to ecdysone*” (GO:0035075) was also enrich in hypomethylated genes in ageing males (Supplemental Figures S2). Genes which are consistently hypermethylated as females age are enriched for “*nuclear-transcribed mRNA catabolic process*” (GO:0000956), see Supplemental Figures S3. Genes which are consistently hypomethylated as females age are enriched for “*negative regulation of JUN kinase activity*” (GO:0043508) and “*protein localization to kinetochore*” (GO:0034501), see Supplemental Figures S4.

### Variably methylated positions

29,012 age-related VMPs were found amongst the 375,403 methylated CpGs. The identified VMPs were located at 2,451 different genes (Supplemental Table S6). Supplemental Figure S5 shows a treemap of GO terms associated with these genes, these include cell surface receptor signalling pathway (GO:0007166), cellular component morphogenesis (GO:0032989), cellular macromolecule localization (GO:0070727) and protein phosphorylation (GO:0006468).

### Epigenetic entropy

There was a significant interaction of chronological age and sex on their effects on epigenetic entropy (Beta regression: χ^2^ = 5.4654, d.f. = 1, p = 0.0019), see Figure 2c. Females display the increasing pattern found in other species (F= 7.491, df = 1, p = 0.0062), but males display no relationship between entropy and chronological age (F= 0.304, df = 1, p = 0.5816).

### Epigenetic clock

The epigenetic clock was constructed by regressing chronological age against the 5290 significantly differentially methylated CpGs. This identified 19 CpGs that best predict age. Eight of these decrease in methylation as *Nasonia* age and eleven increase in methylation. The full list of these CpGs and the genes where they are located can be found in Supplemental Table S7. GO terms associated with these genes can be seen in Supplementary Figure S6. These include glyoxylate cycle (GO:0006097), response to oxidative stress (GO:0006979), induction of programmed cell death (GO:0012502), and regulation of cell growth (GO:0001558).

The epigenetic age of each replicate is the weighted average of these CpGs’ methylation state. This correlates with chronological age (Spearman’s *ρ* = 0.94, p = 1.4 × 10^−8^, see Figure 2d), with a Root Mean Square Error of 0.39 days between chronological age and epigenetic age. Male and females show no difference in age acceleration (Wilcoxon test:W = 35, p = 0.9626), see Supplementary Figure S7.

## Discussion

We have found evidence of dynamic DNA methylation changes associated with ageing in the model insect *Nasonia vitripennis*. This includes differentially methylated CpGs, variably methylated CpGs, an increase in epigenetic entropy in females and an epigenetic clock that accurately predicts age in *Nasonia*. Our methylation data is based on whole body samples. Individual cell or tissue types are known to have larger differences in methylation than are found throughout ageing [49]. Our finding of age-related epigenetic changes despite using whole body samples fits with the use of pan-tissue epigenetic clocks in humans, although see Porter et al.[49].

A limitation of this study is the short time span over which the methylation samples were collected. We based our BS-seq sampling on an earlier study that found that males had an average adult life expectancy of 10.7 days and females 16.6 days [48]. Our wasps lived 67% longer, meaning our methylation analysis captured a reduced amount of the ageing process. Despite this, our results still display clear ageing patterns in methylation. This fits with epigenetic clocks in other animals where there is a linear change in methylation over the course of the adult lifespan [27].

In our survivial experiment, females live longer than males. The survival curves also appear different. Females have the classic type 1 curve, but males appear almost as a type 3 curve with high mortality rates at a younger age, but the rate of death seems to slow down once males go past median lifespan. The increased lifespan of females reflects the pattern found in humans, most mammals and *Drosophila* [50]. In humans, this sex difference in lifespan is universal but in other species, it depends strongly on genotype and environment. Therefore, any general conclusions about sex difference in *Nasonia* ageing will need further experiments using different genotypes and environments.

Overall, DNA methylation increases in *Nasonia* as it ages . This is mirrored in the pattern of differentially methylated genes, with the majority being consistently hypermethylated as *Nasonia* age. This contrasts with the general hypomethylation found in mammalian ageing [51], although this is highly tissue specific with many tissues and cell types showing no relationship between global DNA methylation and ageing [5]. With the proviso that our data comes from whole body analysis, this hypermethylation pattern in ageing *Nasonia* might reflect the different roles of DNA methylation in invertebrates compared to mammals [52].

Females consistently exhibit lower levels of methylation compared to males as they age. Additionally, there is a much lower number of genes showing consistent hypermethylation or hypomethylation in aging females compared to aging males. This trend is reminiscent of a recent finding in the mealybug *Planococcus citri* [53], where males exhibit higher methylation levels distributed across the genome, while females have lower overall methylation but concentrated in specific regions. The authors hypothesize that this pattern is related to the mealybug’s paternal genome elimination process, wherein paternal chromosomes in males are highly condensed and eliminated from sperm. This sex-specific distribution of methylation may also be observed in *Nasonia*. Wang et al. [54] discovered that in *Nasonia*, only a few genes with male-biased expression are methylated, while a considerable number of genes with female-biased expression exhibit methylation. Epigenetic drift, measured as either variably methylated positions or entropy increases as *Nasonia* ages. Nearly 8% of methylated CpGs were classified as variably methylated positions. Amongst them are a number of sirtuins (see Supplementary Table S6), which have been proposed as central to epigenetic ageing [55]. VMPs have been proposed to measure secondary ageing [5]. Primary ageing are changes in all tissues over time, secondary ageing are deleterious changes aggravated by the environment and disease [5]. This idea suggests that VMPs are the part of the methylome that responds to ageing interventions. That is the methylome becomes more youthful by returning to its less noisy original state.

Epigenetic entropy, at least in females, seems to follow the mammalian pattern, with the methylome becoming less predictable as female *Nasonia* age. A recent study showed that epigenetic entropy in a species predicts that species’ maximum lifespan [56]. Males, however show no relationship between chronological age and entropy. We offer two possible explanations of this. Firstly, the male pattern seems to be reflected in the survivourship curve (Figure 2a), where our day 16 samples almost correspond to mean male lifespan (17 days). After this point, the rate of death seems to slow down. This slower rate of death might be associated with this decreased entropy in sixteen day old males, akin to the late life cessation of age-related deterioration found in many species [57]. Secondly, the lifespan of males appears more variable than females. This is most clearly seen in the age acceleration of the two sexes, see Supplementary Figure S7. This increased variation in male lifespan could mean, with our small sample size, it is more difficult to find the relationship with entropy in males.

The tight correlation between chronological age and epigenetic age as calculated by the epigenetic clock is to be expected. There is very little variation in our test subjects. They all come from a single strain and are kept in exactly the same environment. That is, there is nothing to induce variation in biological ageing. We also found no age acceleration between males and females. This epigenetic clock is similar to results in many vertebrates [58] and even recently in the crustacean, the water flea *Daphnia magna* [9]. However, this is the first time an epigenetic clock has been discovered in a tractable insect model.

Unsurprisingly, there is no direct correspondence between the thirteen genes found in multiple human epigenetic clocks [59] and the genes associated with our nineteen clock CpGs. Although, the CpG having the most effect on epigenetic age in *Nasonia* is located in the gene for a leucine-rich repeat kinase (lrrk). LRRK2 mutations are a common cause of age related autosomal-dominant Parkinson’s disease [60]. Genes associated with neurodegenerative diseases are common in mammalian epigenetic clocks. There is some overlap in the GO terms associated with a pan-mammalian epigenetic clock [61] and those we found in *Nasonia* (Supplementary Figure S6). These include nucleic acid binding and RNA metabolic processes.

We predict two main areas where our establishment of an epigenetic clock in a model insect species will be useful; firstly, the biology underpinning epigenetic clocks and secondly, how influenced epigenetic clocks are by ageing interventions. Variation in the rate of an individual’s epigenetic clock is affected by a large number of traits including inflammation, cell division, metabolic effects, cellular heterogeneity, diet, and numerous other lifestyle factors [26]. *Nasonia*, with its simplified insect systems, is perfect to experimentally separate the different processes involved in the biology of the clock into its constitutive parts [12]. Being short-lived (3-4 weeks as opposed to 26-30 months for mice), *Nasonia* are ideal to measure the effects of ageing interventions on both life span and epigenetic ageing. This will answer the question does a short-term decrease in someone’s epigenetic clock score lower their chance of developing age-related ill health, that is if epigenetic clocks can be used as endpoints for clinical trials of various anti-ageing interventions [28].

Starting with Medawar, genetic mutations were seen as the driver of ageing [62]. Recent theories on the causes of ageing focus rather on the loss of epigenetic information as the main driver of ageing [55]. These epigenetic factors due to their known plasticity, are tempting targets for anti-ageing interventions [63]. We propose *Nasonia vitripennis*, with its fully functional DNA methylation system and its now established ageing methylome as a model for this epigenetic era of ageing research.

## Acknowledgements

EBM was funded by grant RPG-2020-363 from the Leverhulme Trust. CT, KB, BC, TWC and ARCJ were supported by a BBSRC MIBTP DTP studentships. EBM and CT were funded by pioneer grant APP3335 from the BBSRC. This research used the ALICE2 High Performance Computing Facility at the University of Leicester.

## Data accessibility

All sequencing data related to this project can be found under EBI ArrayExpress E-MTAB-13854. Custom scripts for the genome-wide analysis can be found at https://github.com/EamonnMallon/nasonia_epigenetic_ageing.

## Authors’ contributions

EBM designed the study. CT and KB carried out husbandry, sampling and DNA extraction. EBM, CT, ARCJ, KB and BC analysed the data. EBM wrote the initial draft. All authors were involved in redrafting.

## Supplementary figures and text

**Figure S1:**
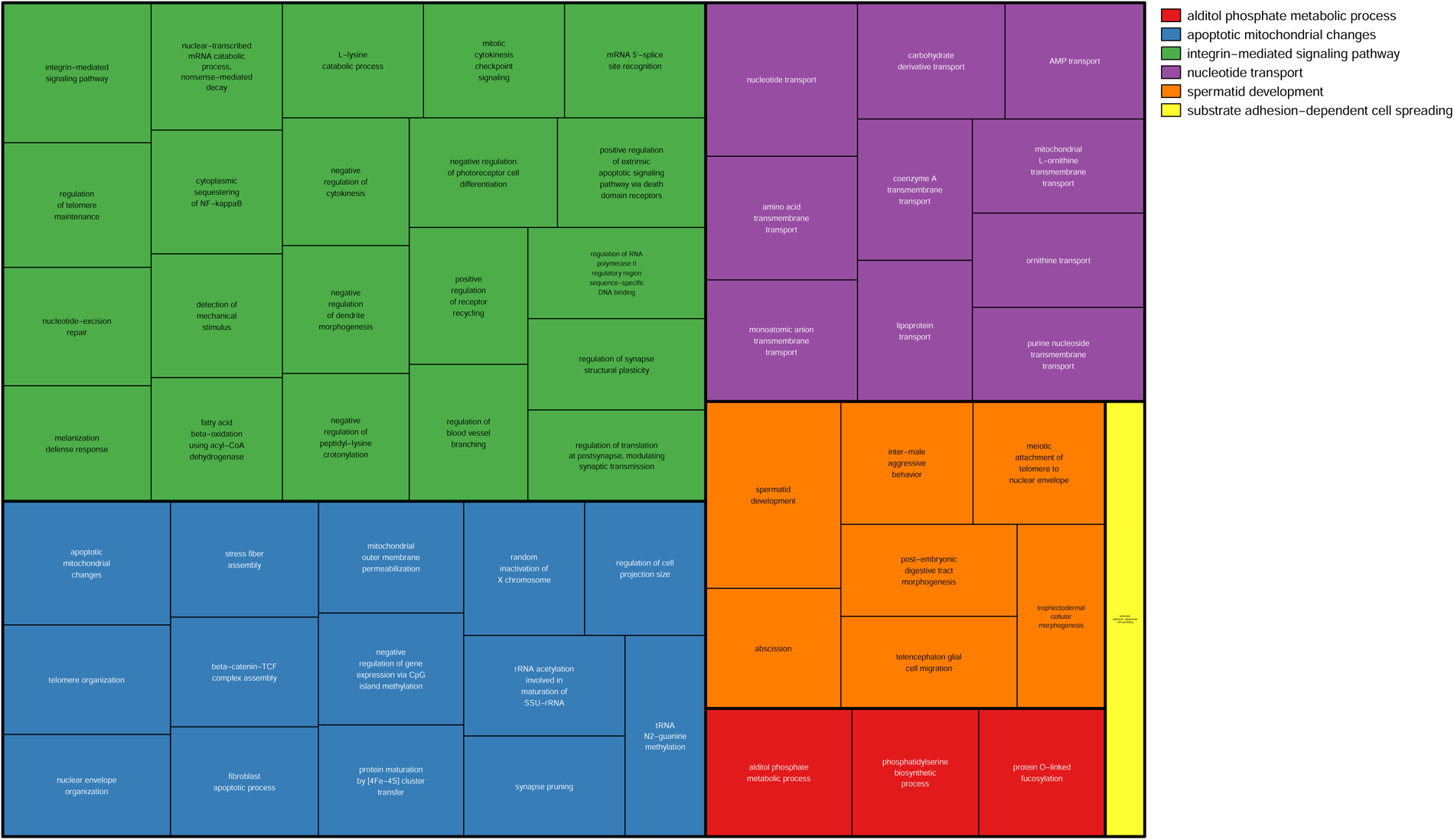
Treemap of Biological Processes GO term enrichment for consistently hypermethylated genes associated with ageing in males. Enriched BP for GO terms (p <0.05) clustered using REVIGO. These rectangles are joined into different coloured ‘superclusters’ of loosely related terms. The area of the rectangles represents the p-value associated with that cluster’s enrichment.

**Figure S2:**
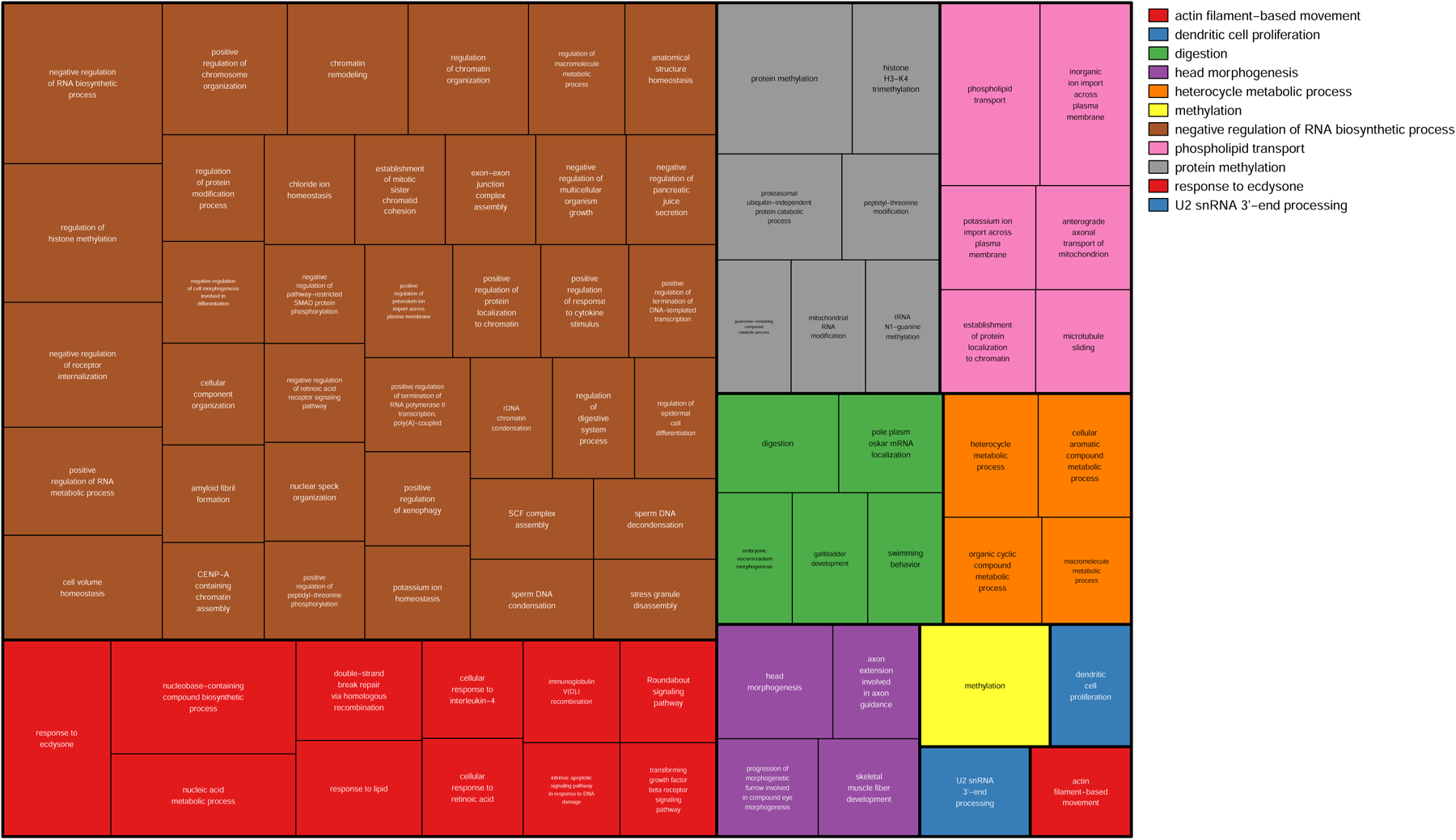
Treemap of Biological Processes GO term enrichment for consistently hypomethylated genes associated with ageing in males. Enriched BP for GO terms (p <0.05) clustered using REVIGO. These rectangles are joined into different coloured ‘superclusters’ of loosely related terms. The area of the rectangles represents the p-value associated with that cluster’s enrichment.

**Figure S3:**
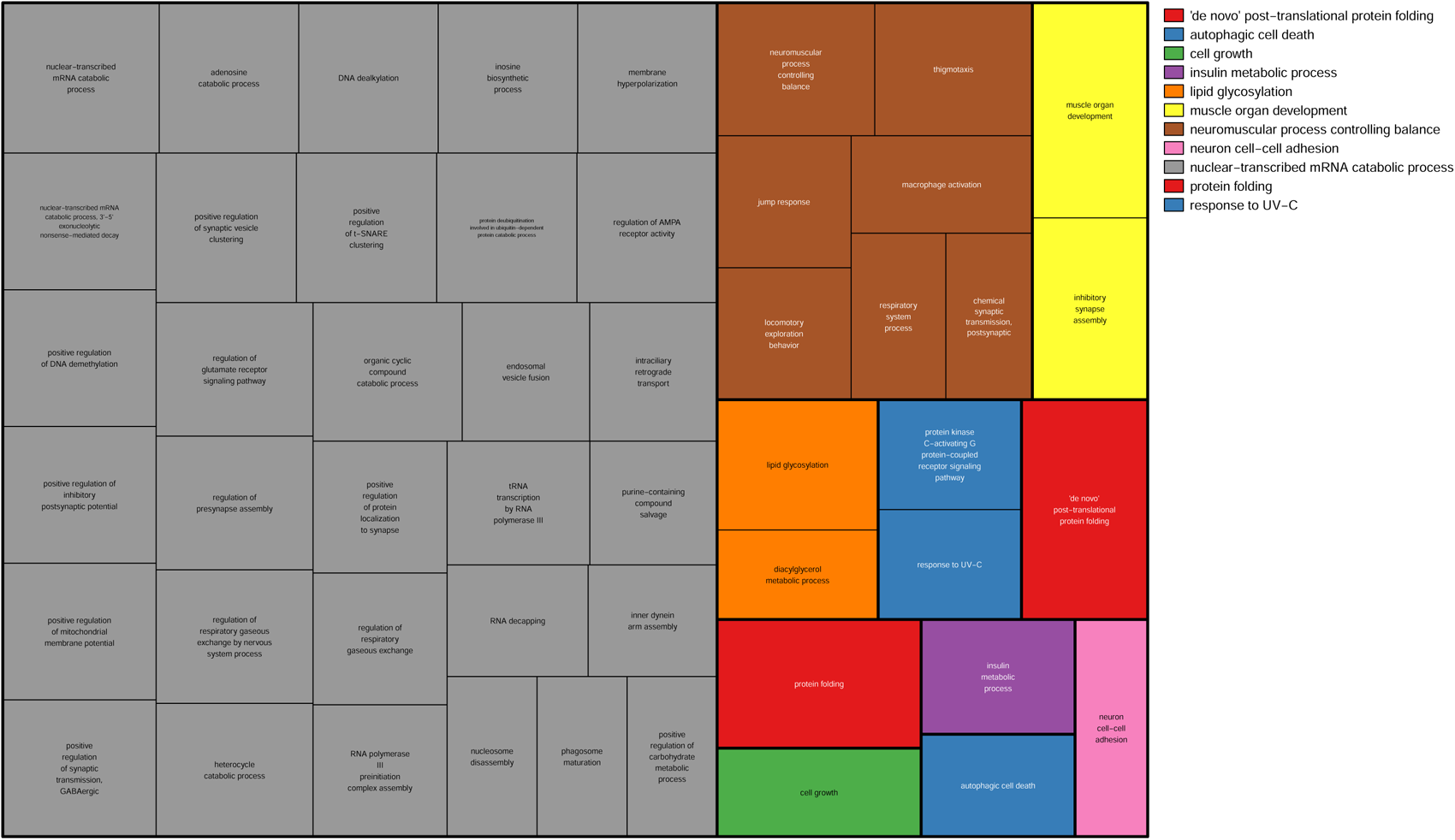
Treemap of Biological Processes GO term enrichment for consistently hypermethylated genes associated with ageing in females. Enriched BP for GO terms (p <0.05) clustered using REVIGO. These rectangles are joined into different coloured ‘superclusters’ of loosely related terms. The area of the rectangles represents the p-value associated with that cluster’s enrichment.

**Figure S4:**
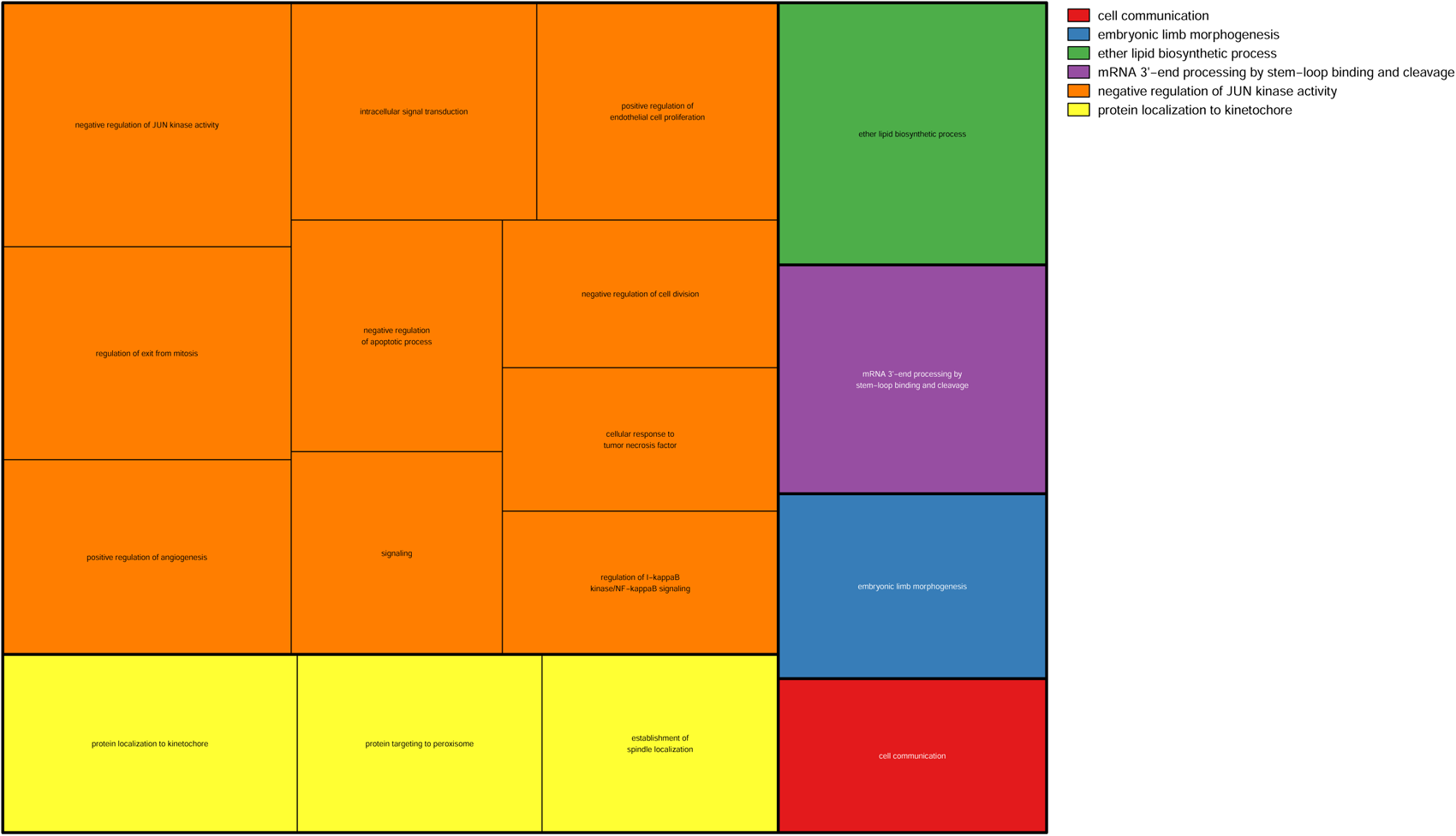
Treemap of Biological Processes GO term enrichment for consistently hypomethylated genes associated with ageing in females. Enriched BP for GO terms (p <0.05) clustered using REVIGO. These rectangles are joined into different coloured ‘superclusters’ of loosely related terms. The area of the rectangles represents the p-value associated with that cluster’s enrichment.

**Figure S5:**
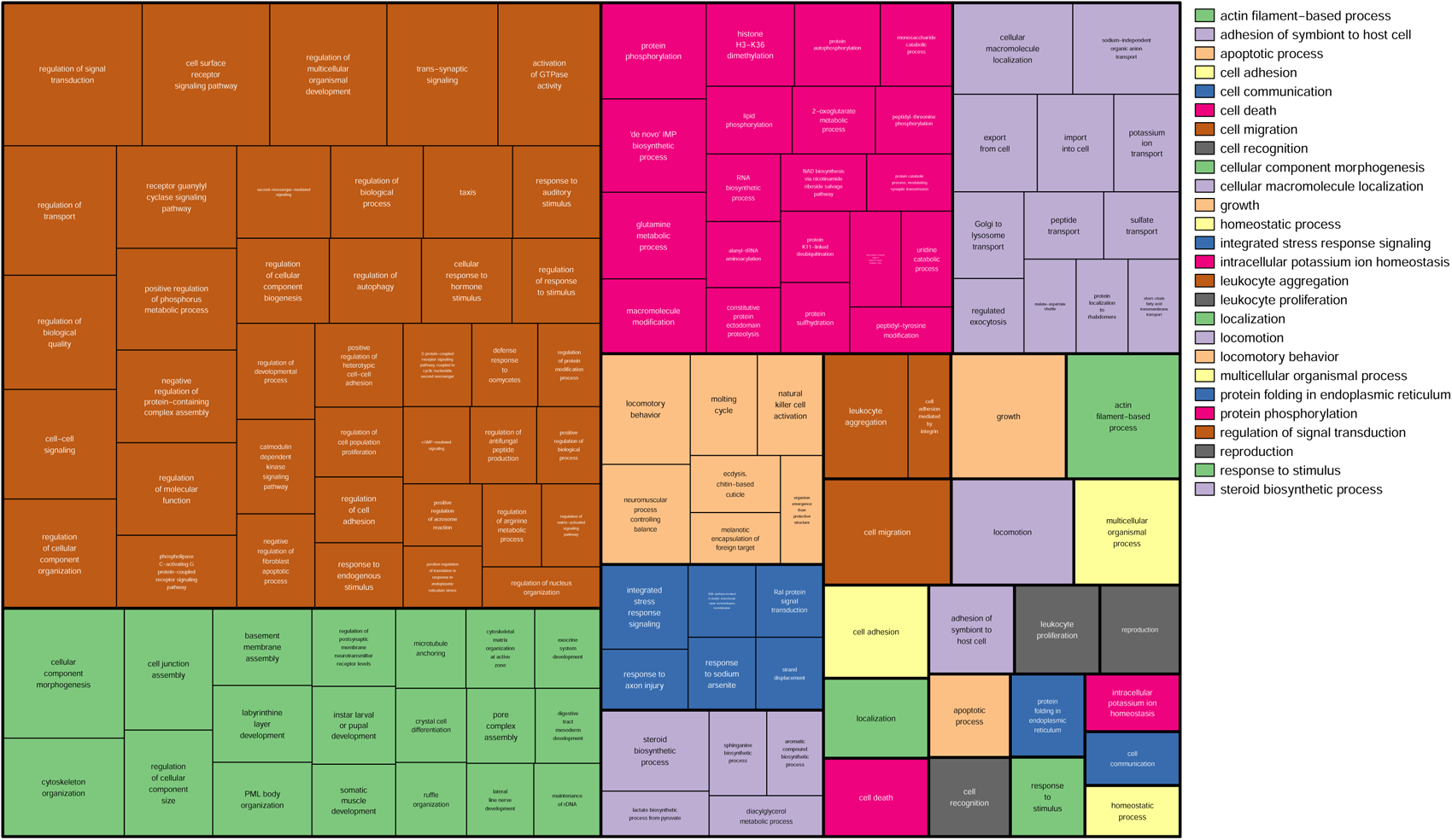
Treemap of Biological Processes GO term enrichment for variably methylated positions associated with ageing. Enriched BP for GO terms (p <0.05) clustered using REVIGO. These rectangles are joined into different coloured ‘superclusters’ of loosely related terms. The area of the rectangles represents the p-value associated with that cluster’s enrichment.

**Figure S6:**
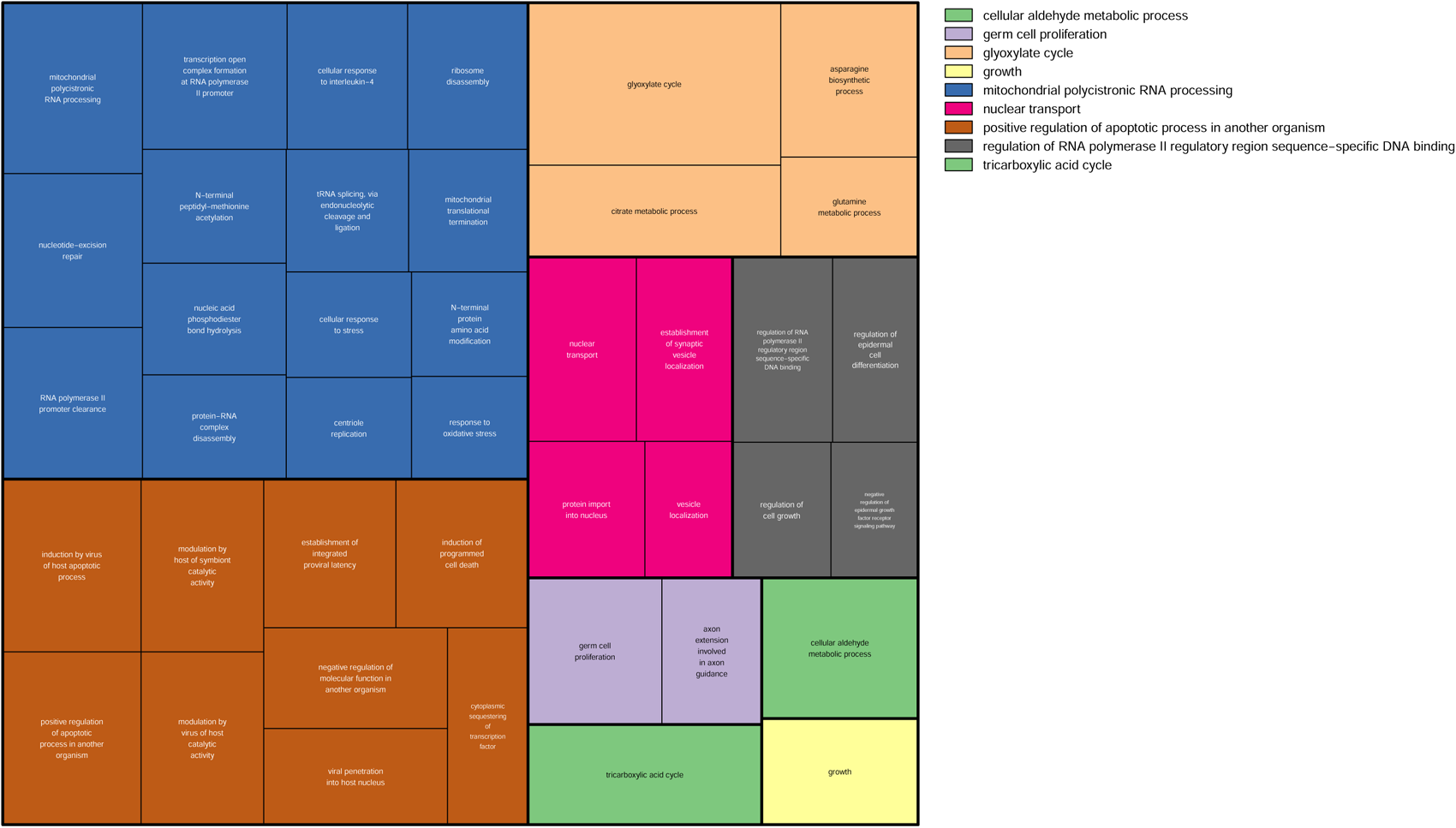
Treemap of Biological Processes GO term enrichment for epigenetic clock genes associated. Enriched BP for GO terms (p <0.05) clustered using REVIGO. These rectangles are joined into different coloured ‘superclusters’ of loosely related terms. The area of the rectangles represents the p-value associated with that cluster’s enrichment.

**Figure S7:**
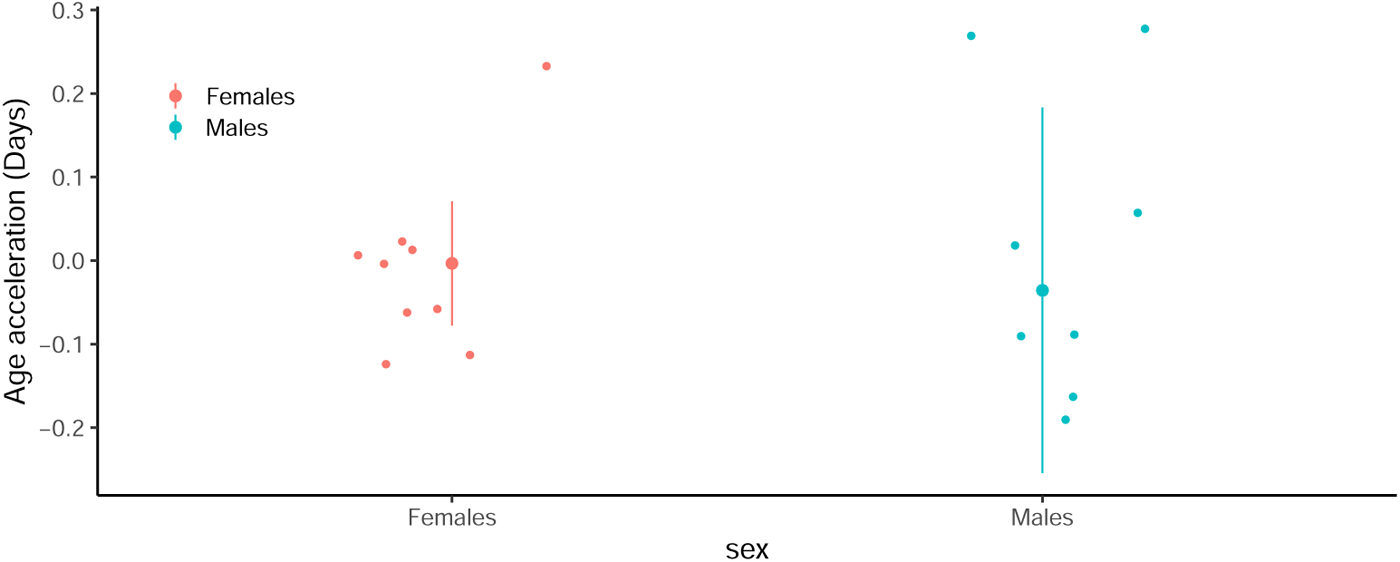
Age acceleration calculated as the residual of chronological age against epigenetic age for each sample. The large central dots represent the median value, with the vertical lines representing interquartile range.

## References

[1] Hildegard I. D. Mack, Thomas Heimbucher, and Coleen T. Murphy. The nematode Caenorhabditis elegans as a model for aging research. Drug Discovery Today: Disease Models, 27:3–13, March 2018. ISSN 1740-6757. doi: 10.1016/j.ddmod.2018.11.001. URL https://www.sciencedirect.com/science/article/pii/S1740675718300343.

[2] Matthew D. W. Piper and Linda Partridge. Drosophila as a model for ageing. Biochimica et Biophysica Acta (BBA) - Molecular Basis of Disease, 1864(9, Part A):2707–2717, September 2018. ISSN 0925-4439. doi: 10.1016/j.bbadis.2017.09.016. URL https://www.sciencedirect.com/science/article/pii/S0925443917303319.

[3] Frank Lyko and Ryszard Maleszka. Insects as Innovative Models for Functional Studies of DNA Methylation. Trends in genetics, 27(4):127–131, April 2011. ISSN 0168-9525. doi: 10.1016/j.tig.2011.01.003.

[4] Chiung-Wen Hu, Jian-Lian Chen, Yu-Wen Hsu, Cheng-Chieh Yen, and Mu-Rong Chao. Trace analysis of methylated and hydroxymethylated cytosines in DNA by isotope-dilution LC-MS/MS: first evidence of DNA methylation in Caenorhabditis elegans. The Biochemical Journal, 465(1):39–47, January 2015. ISSN 1470-8728. doi: 10.1042/BJ20140844.

[5] Kirsten Seale, Steve Horvath, Andrew Teschendorff, Nir Eynon, and Sarah Voisin. Making Sense of the Ageing Methylome. Nature Reviews Genetics, pages 1–21, May 2022. ISSN 1471-0064. doi: 10.1038/s41576-022-00477-6. URL https://www.nature.com/articles/s41576-022-00477-6. Publisher: Nature Publishing Group.

[6] Claire Morandin, Volker P. Brendel, Liselotte Sundström, Heikki Helanterä, and Alexander S. Mikheyev. Changes in gene DNA methylation and expression networks accompany caste specialization and age-related physiological changes in a social insect. Molecular Ecology, 28(8):1975–1993, 2019. ISSN 1365-294X. doi: 10.1111/mec.15062. URL http://onlinelibrary.wiley.com/doi/abs/10.1111/mec.15062. eprint: https://onlinelibrary.wiley.com/doi/pdf/10.1111/mec.15062.

[7] Thibaut Renard, Baptiste Martinet, Natalia De Souza Araujo, and Serge Aron. DNA methylation extends lifespan in the bumblebee Bombus terrestris. Proceedings of the Royal Society B: Biological Sciences, 290(2012):20232093, December 2023. doi: 10.1098/rspb.2023.2093. URL https://royalsocietypublishing.org/doi/full/10.1098/rspb.2023.2093. Publisher: Royal Society.

[8] Carlos A. M. Cardoso-Júnior, Karina R. Guidugli-Lazzarini, and Klaus Hartfelder. DNA methylation affects the lifespan of honey bee (Apis mellifera L.) workers - Evidence for a regulatory module that involves vitellogenin expression but is independent of juvenile hormone function. Insect Biochemistry and Molecular Biology, 92:21–29, January 2018. ISSN 1879-0240. doi: 10.1016/j.ibmb.2017.11.005.

[9] Jack Hearn, Fiona Plenderleith, and Tom J. Little. DNA methylation differs extensively between strains of the same geographical origin and changes with age in Daphnia magna. Epigenetics & Chromatin, 14(1):4, January 2021. ISSN 1756-8935. doi: 10.1186/s13072-020-00379-z. URL https://doi.org/10.1186/s13072-020-00379-z.

[10] Akiko Koto, Makoto Tamura, Pui Shan Wong, et al. Social isolation shortens lifespan through oxidative stress in ants. Nature Communications, 14(1):5493, September 2023. ISSN 2041-1723. doi: 10.1038/s41467-023-41140-w. URL https://www.nature.com/articles/s41467-023-41140-w. Number: 1 Publisher: Nature Publishing Group.

[11] Benjamin P. Oldroyd and Boris Yagound. The role of epigenetics, particularly DNA methylation, in the evolution of caste in insect societies. Philosophical Transactions of the Royal Society B: Biological Sciences, 376(1826):20200115, 2020. ISSN 0962-8436. doi: 10.1098/rstb.2020. 0115. URL https://www.ncbi.nlm.nih.gov/pmc/articles/PMC8059649/.

[12] John H. Werren, Stephen Richards, Christopher A. Desjardins, et al. Functional and evolutionary insights from the genomes of three parasitoid Nasonia species. Science (New York, N.Y.), 327 (5963):343–348, January 2010. ISSN 1095-9203. doi: 10.1126/science.1178028.

[13] Xu Wang, David Wheeler, Amanda Avery, et al. Function and Evolution of DNA Methylation in Nasonia vitripennis. PLOS Genetics, 9(10):e1003872, October 2013. ISSN 1553-7404. doi: 10.1371/journal.pgen.1003872. URL http://journals.plos.org/plosgenetics/article?id=10.1371/journal.pgen.1003872.

[14] Suzannah M Beeler, Garrett T Wong, Jennifer M Zheng, et al. Whole-genome DNA methylation profile of the jewel wasp (Nasonia vitripennis). G3: Genes, Genomes, Genetics, 4(3):383–388, 2014.

[15] Deanna Arsala, Xin Wu, Soojin V. Yi, and Jeremy A. Lynch. Dnmt1a is essential for gene body methylation and the regulation of the zygotic genome in a wasp. PLOS Genetics, 18(5):e1010181, May 2022. ISSN 1553-7404. doi: 10.1371/journal.pgen.1010181. URL https://journals.plos.org/plosgenetics/article?id=10.1371/journal.pgen.1010181. Publisher: Public Library of Science.

[16] Mirko Pegoraro, Akanksha Bafna, Nathaniel J. Davies, David M. Shuker, and Eran Tauber. DNA methylation changes induced by long and short photoperiods in Nasonia. Genome Research, 26(2):203–210, February 2016. ISSN 1549-5469. doi: 10.1101/gr.196204.115.

[17] Nicola Cook, Bart A. Pannebakker, Eran Tauber, and David M. Shuker. DNA Methylation and Sex Allocation in the Parasitoid Wasp Nasonia vitripennis. The American Naturalist, 186(4):513–518, October 2015. ISSN 0003-0147. doi: 10.1086/682950. URL https://www.journals.uchicago.edu/doi/full/10.1086/682950. Publisher: The University of Chicago Press.

[18] Marie-Theres Multerer, Martina Wendler, and Joachim Ruther. The biological significance of lipogenesis in Nasonia vitripennis. *Proceedings*. Biological Sciences, 289(1972):20220208, April 2022. ISSN 1471-2954. doi: 10.1098/rspb.2022.0208.

[19] Loan Davies. A Study of the Effect of Diet on the Life-Span of Nasonia Vitripennis (Walk.) (Hymenoptera, Pteromalidae)1. Journal of Gerontology, 30(3):294–298, May 1975. ISSN 0022-1422. doi: 10.1093/geronj/30.3.294. URL https://doi.org/10.1093/geronj/30.3.294.

[20] Kelley Leung, Louis van de Zande, and Leo W. Beukeboom. Life-history traits of the Whiting polyploid line of the parasitoid Nasonia vitripennis. Entomologia Experimentalis et Applicata, 167(7):655–669, 2019. ISSN 1570-7458. doi: 10.1111/eea.12808. URL https://onlinelibrary.wiley.com/doi/abs/10.1111/eea.12808. eprint: https://onlinelibrary.wiley.com/doi/pdf/10.1111/eea.12808.

[21] D. S. Saunders. The effect of the age of female *Nasonia vitripennis* (Walker) (Hymenoptera, Pteromalidae) upon the incidence of larval diapause. Journal of Insect Physiology, 8(3): 309–318, May 1962. ISSN 0022-1910. doi: 10.1016/0022-1910(62)90034-3. https://www.sciencedirect.com/science/article/pii/0022191062900343.

[22] G. J. C. Smith and David Pimentel. The Effect of Two Host Species on the Longevity and Fertility of Nasonia vitripennis1,2. Annals of the Entomological Society of America, 62 (2):305–308, March 1969. ISSN 0013-8746. doi: 10.1093/aesa/62.2.305. URL https://doi.org/10.1093/aesa/62.2.305.

[23] Felix Krueger, Benjamin Kreck, Andre Franke, and Simon R Andrews. DNA methylome analysis using short bisulfite sequencing data. Nature methods, 9(2):145, 2012.

[24] Alexander Vaiserman. Developmental Tuning of Epigenetic Clock. Frontiers in Genetics, 9, 2018. ISSN 1664-8021. URL https://www.frontiersin.org/articles/10.3389/fgene.2018.00584.

[25] Gregory Hannum, Justin Guinney, Ling Zhao, et al. Genome-wide Methylation Profiles Reveal Quantitative Views of Human Aging Rates. Molecular Cell, 49(2):359–367, January 2013. ISSN 1097-2765. doi: 10.1016/j.molcel.2012.10.016. URL https://www.sciencedirect.com/science/article/pii/S1097276512008933.

[26] Christopher G. Bell, Robert Lowe, Peter D. Adams, et al. DNA methylation aging clocks: challenges and recommendations. Genome Biology, 20(1):249, November 2019. ISSN 1474-760X. doi: 10.1186/s13059-019-1824-y. URL https://doi.org/10.1186/s13059-019-1824-y.

[27] Steve Horvath. DNA methylation age of human tissues and cell types. Genome Biology, 14(10):3156, December 2013. ISSN 1474-760X. doi: 10.1186/gb-2013-14-10-r115. URL https://doi.org/10.1186/gb-2013-14-10-r115.

[28] Liam Drew. Turning back time with epigenetic clocks. Nature, 601(7893):S20–S22, January 2022. doi: 10.1038/d41586-022-00077-8. URL https://www.nature.com/articles/d41586-022-00077-8. Bandiera abtest: a Cg type: Outlook Number: 7893 Publisher: Nature Publishing Group Subject term: Ageing, Society, Epigenetics.

[29] Terry M Therneau. A package for survival analysis in R. manual, 2022. URL https://CRAN.R-project.org/package=survival.

[30] Terry M. Therneau. coxme: Mixed effects cox models. manual, 2022. URL https://CRAN.R-project.org/package=coxme.

[31] R Core Team. R: A language and environment for statistical computing. manual, Vienna, Austria, 2021. URL https://www.R-project.org/. tex.organization: R Foundation for Statistical Computing.

[32] S. Andrews. FastQC: a quality control tool for high throughput sequence data, 2010. URL http://www.bioinformatics.babraham.ac.uk/projects/fastqc.

[33] Elena Dalla Benetta, Igor Antoshechkin, Ting Yang, et al. Genome elimination mediated by gene expression from a selfish chromosome. Science Advances, 6(14):eaaz9808, April 2020. doi: 10.1126/sciadv.aaz9808. URL https://www.science.org/doi/10.1126/sciadv.aaz9808. Publisher: American Association for the Advancement of Science.

[34] Ben Langmead and Steven L Salzberg. Fast gapped-read alignment with Bowtie 2. Nature methods, 9(4):357–359, March 2012. ISSN 1548-7091. doi: 10.1038/nmeth.1923. URL https://www.ncbi.nlm.nih.gov/pmc/articles/PMC3322381/.

[35] Felix Krueger and Simon R. Andrews. Bismark: a flexible aligner and methylation caller for Bisulfite-Seq applications. Bioinformatics, 27(11):1571–1572, June 2011. ISSN 1367-4803. doi: 10.1093/bioinformatics/btr167. URL https://academic.oup.com/bioinformatics/article/27/11/1571/216956.

[36] Heng Li, Bob Handsaker, Alec Wysoker, et al. The Sequence Alignment/Map format and SAMtools. Bioinformatics (Oxford, England), 25(16):2078–2079, August 2009. ISSN 1367-4811. doi: 10.1093/bioinformatics/btp352.

[37] Altuna Akalin, Matthias Kormaksson, Sheng Li, et al. methylKit: a comprehensive R package for the analysis of genome-wide DNA methylation profiles. Genome Biology, 13(10):R87, October 2012. ISSN 1474-760X. doi: 10.1186/gb-2012-13-10-r87.

[38] Longjie Cheng and Yu Zhu. A classification approach for DNA methylation profiling with bisulfite next-generation sequencing data. *Bioinformatics (Oxford*, England), 30(2):172–179, January 2014. ISSN 1367-4811. doi: 10.1093/bioinformatics/btt674.

[39] Matthew D. Schultz, Robert J. Schmitz, and Joseph R. Ecker. ‘Leveling’ the playing field for analyses of single-base resolution DNA methylomes. Trends in genetics : TIG, 28 (12):583–585, December 2012. ISSN 0168-9525. doi: 10.1016/j.tig.2012.10.012. URL https://www.ncbi.nlm.nih.gov/pmc/articles/PMC3523709/.

[40] Bettina Grün, Ioannis Kosmidis, and Achim Zeileis. Extended Beta Regression in R: Shaken, Stirred, Mixed, and Partitioned. Journal of Statistical Software, 48:1–25, May 2012. ISSN 1548-7660. doi: 10.18637/jss.v048.i11. URL https://doi.org/10.18637/jss.v048.i11.

[41] Achim Zeileis and Torsten Hothorn. Diagnostic Checking in Regression Relationships. 2, 2002.

[42] Francisco Cribari-Neto and Achim Zeileis. Beta Regression in R. Journal of Statistical Software, 34:1–24, April 2010. ISSN 1548-7660. doi: 10.18637/jss.v034.i02. URL https://doi.org/10.18637/jss.v034.i02.

[43] Russell V. Lenth. emmeans: Estimated marginal means, aka least-squares means. manual, 2022. URL https://CRAN.R-project.org/package=emmeans.

[44] Jerome Friedman, Trevor Hastie, and Robert Tibshirani. Regularization paths for generalized linear models via coordinate descent. Journal of Statistical Software, 33(1):1–22, 2010. doi: 10.18637/jss.v033.i01. URL https://www.jstatsoft.org/v33/i01/.

[45] Fiona Cunningham, James E Allen, Jamie Allen, et al. Ensembl 2022. Nucleic Acids Research, 50(D1):D988–D995, January 2022. ISSN 0305-1048. doi: 10.1093/nar/gkab1049. URL https://doi.org/10.1093/nar/gkab1049.

[46] Yoav Benjamini and Yosef Hochberg. Controlling the False Discovery Rate: A Practical and Powerful Approach to Multiple Testing. Journal of the Royal Statistical Society. Series B (Methodological), 57(1):289–300, 1995. ISSN 0035-9246. URL http://www.jstor.org/stable/2346101.

[47] Fran Supek, Matko Bošnjak, Nives Škunca, and Tomislav Šmuc. REVIGO Summarizes and Visualizes Long Lists of Gene Ontology Terms. PLOS ONE, 6(7):e21800, July 2011. ISSN 1932-6203. doi: 10.1371/journal.pone.0021800. URL http://journals.plos.org/plosone/article?id=10.1371/journal.pone.0021800.

[48] Theresa S. E. Floessner, Floor E. Boekelman, Stella J. M. Druiven, et al. Lifespan is unaffected by size and direction of daily phase shifts in Nasonia, a hymenopteran insect with strong circadian light resetting. Journal of Insect Physiology, 117:103896, September 2019. ISSN 1879-1611. doi: 10.1016/j.jinsphys.2019.103896.

[49] Hunter L. Porter, Chase A. Brown, Xiavan Roopnarinesingh, et al. Many chronological aging clocks can be found throughout the epigenome: Implications for quantifying biological aging. Aging Cell, 20(11):e13492, November 2021. ISSN 1474-9718. doi: 10.1111/acel.13492. URL https://www.ncbi.nlm.nih.gov/pmc/articles/PMC8590098/.

[50] Steven N. Austad and Kathleen E. Fischer. Sex Differences in Lifespan. Cell metabolism, 23(6):1022–1033, June 2016. ISSN 1550-4131. doi: 10.1016/j.cmet.2016.05.019. URL https://www.ncbi.nlm.nih.gov/pmc/articles/PMC4932837/.

[51] Marc Jung and Gerd P. Pfeifer. Aging and DNA methylation. BMC Biology, 13(1):7, January 2015. ISSN 1741-7007. doi: 10.1186/s12915-015-0118-4. URL https://doi.org/10.1186/s12915-015-0118-4.

[52] H. Marshall, M. T. Nicholas, J. S. van Zweden, et al. DNA methylation is associated with codon degeneracy in a species of bumblebee. Heredity, 130(4):188–195, April 2023. ISSN 1365-2540. doi: 10.1038/s41437-023-00591-z. URL https://www.nature.com/articles/s41437-023-00591-z. Number: 4 Publisher: Nature Publishing Group.

[53] Stevie A. Bain, Hollie Marshall, Andrés G. de la Filia, et al. Sex-specific expression and DNA methylation in a species with extreme sexual dimorphism and paternal genome elimination. Molecular Ecology, 30(22):5687–5703, November 2021. ISSN 1365-294X. doi: 10.1111/mec.15842.

[54] Xu Wang, John H. Werren, and Andrew G. Clark. Genetic and epigenetic architecture of sex-biased expression in the jewel wasps Nasonia vitripennis and giraulti. Proceedings of the National Academy of Sciences of the United States of America, 112(27):E3545–E3554, July 2015. ISSN 0027-8424. doi: 10.1073/pnas.1510338112. URL https://www.ncbi.nlm.nih.gov/pmc/articles/PMC4500217/.

[55] Jae-Hyun Yang, Motoshi Hayano, Patrick T. Griffin, et al. Loss of epigenetic information as a cause of mammalian aging. Cell, 186:1–22, January 2023. ISSN 0092-8674, 1097-4172. doi: 10.1016/j.cell.2022.12.027. URL https://www.cell.com/cell/abstract/S0092-8674(22)01570-7. Publisher: Elsevier.

[56] Emily M. Bertucci-Richter and Benjamin B. Parrott. The rate of epigenetic drift scales with maximum lifespan across mammals. Nature Communications, 14(1):7731, November 2023. ISSN 2041-1723. doi: 10.1038/s41467-023-43417-6. URL https://www.nature.com/articles/s41467-023-43417-6. Number: 1 Publisher: Nature Publishing Group.

[57] Michael R. Rose, Casandra L. Rauser, and Laurence D. Mueller. Late Life: A New Frontier for Physiology. Physiological and Biochemical Zoology, 78(6):869–878, November 2005. ISSN 1522-2152. doi: 10.1086/498179. URL https://www.journals.uchicago.edu/doi/abs/10.1086/498179. Publisher: The University of Chicago Press.

[58] Benjamin B. Parrott and Emily M. Bertucci. Epigenetic Aging Clocks in Ecology and Evolution. Trends in Ecology & Evolution, 34(9):767–770, September 2019. ISSN 1872-8383. doi: 10.1016/j.tree.2019.06.008.

[59] Adam Li, Zane Koch, and Trey Ideker. Epigenetic aging: Biological age prediction and informing a mechanistic theory of aging. Journal of Internal Medicine, 292(5):733–744, 2022. ISSN 1365-2796. doi: 10.1111/joim.13533. URL https://onlinelibrary.wiley.com/doi/abs/10.1111/joim.13533. eprint: https://onlinelibrary.wiley.com/doi/pdf/10.1111/joim.13533.

[60] Jinhua Chen, Ying Chen, and Jiali Pu. Leucine-Rich Repeat Kinase 2 in Parkinson’s Disease: Updated from Pathogenesis to Potential Therapeutic Target. European Neurology, 79(5-6):256–265, 2018. ISSN 0014-3022, 1421-9913. doi: 10.1159/000488938. URL https://www.karger.com/Article/FullText/488938. Publisher: Karger Publishers.

[61] A. T. Lu, Z. Fei, A. Haghani, et al. Universal DNA methylation age across mammalian tissues. Nature Aging, 3(9):1144–1166, September 2023. ISSN 2662-8465. doi: 10.1038/ s43587-023-00462-6.

[62] Leo Szilard. On the nature of the aging process. Proceedings of the National Academy of Sciences, 45(1):30–45, January 1959. doi: 10.1073/pnas.45.1.30. URL https://www.pnas.org/doi/abs/10.1073/pnas.45.1.30. Publisher: Proceedings of the National Academy of Sciences.

[63] Noémie Gensous, Claudio Franceschi, Aurelia Santoro, et al. The Impact of Caloric Restriction on the Epigenetic Signatures of Aging. International Journal of Molecular Sciences, 20(8): 2022, April 2019. ISSN 1422-0067. doi: 10.3390/ijms20082022. URL https://www.ncbi.nlm.nih.gov/pmc/articles/PMC6515465/.

